# Removal Of Environmental Pollutants (Lead, Chromium And Cadmium) Using Root And Leaf Tissues Of Indian Mustard, Rice And Wheat Plants

**DOI:** 10.1101/2022.02.08.479640

**Authors:** Shema Halder, Apurba Anirban

## Abstract

The environment is polluted day by day and it is very much expensive to remediate the environmental pollutants by physicochemical process. Therefore, using plants as a process to remediate pollutants is essential. In this study, phytoremediation of root and leaf tissues of Indian mustard (*Brassica juncea*), rice (*Oryza sativa*) and wheat (*Triticum aestivum*) against three most environment pollutants *viz*. lead (Pb), chromium (Cr) and cadmium (Cd) of Buriganga riverbank soil of Dhaka city, Bangladesh were assessed. The highest amount of Pb was found in the leaf (11.6755 ±1.9860 mg/kg dry weight) and root (51.4251 ±5.0320 mg/kg dry weight) of *B. juncea*. However, the highest amount of Cr was found in the leaf (5.9871±0.9032 mg/kg dry weight) of *B. juncea*; and in the root (46.4739±2.2920 mg/kg dry weight) of *O. sativa* respectively. Although the highest amount (0.9624±0.0920 mg/kg dry weight) of Cd was found in the leaf of *B. juncea*; the amount of Cd in the root was approximately same in all the three plants. This research also found more amounts of heavy metals compared to other studies using Indian mustard, rice and wheat against the three most environmental pollutants viz., Pb, Cr and Cd. Moreover, this study revealed that *B. juncea* is the highest hyperaccumulator species regarding Pb, Cr and Cd accumulation and can be used to clean up the polluted Buriganga riverbank soil.

## INTRODUCTION

Due to the increase of population Industrialization as well as urbanization is taking place simultaneously. As a result, to mitigate the need of everyday life, various industries are producing different commercial products, which are also responsible for producing different waste materials, pollutants, and heavy metals. These contaminants are migrating to the environment and creating contamination of the ecosystems (Gaur, 2004). Dhaka city disposes huge amount of solid waste every day and most of it is released into Buriganga river. Buriganga is the economically very important river of Dhaka city and is polluted by chemical waste, medical waste, household waste, transportation waste, sewage, dead animals, plastic and many other pollutants. These wastes possess different heavy metals, which are responsible for harmful effect on the adjacent aquatic plants, animals as well as on human health. Moreover, heavy metal is responsible for toxic effects (Shukla and Singhal, 1984) as well as oxidative stress (Ahmad et al., 2011) to plants. Heavy metal toxicity also has impact on nutrition and metabolism of plants (John et al., 2009).

Phytoremediation is regarded as the removal of hazardous contaminants from air, water, and soil by using different types of plants (Prasad and Freitas, 2003). It is one of the best methods to remediate pollutants of contaminated groundwater and surface soil (Mwegoha, 2008), which are removed with different costly methods *viz*. physical, chemical, and thermal processes. It was estimated that removing one metre^3^ soil from a polluted site is in the range of USD 0.6–2.5 million (McIntyre, 2003). In contrast, phytoremediation is a cheaper, eco-friendly, as well as alternative to traditional methods. The cost of phytoremediation is $37.7/m^3^ when used plants and trees which have deep roots (Wan et al., 2016). As such, phytoremediation is an innovative cheaper technology than traditional technologies to combat with pollutants (Garbisu and Alkorta, 2001; McIntyre and Lewis, 1997).

To know which plants are suitable to be considered as hyperaccumulators is important for the study of phytoremediation (Rodriguez et al., 2005). By several investigations it was proved that Indian mustard (Choudhury et al., 2016; Ishikawa et al., 2006; John et al., 2009; Salido et al., 2003), rice (Hussain et al., 2021; Murakami et al., 2009; Payus and Talip, 2014; Satpathy et al., 2014; Zakaria et al., 2021), wheat (Chandra et al., 2009; Liu et al., 2009), maize (Wang et al., 2002) etc. are some plants which were capable of eliminating heavy metals. Moreover, other plants were also used to remove different contaminants (Ampiah-Bonney et al., 2007; Maine et al., 2021; Melo et al., 2021; Pratas et al., 2012).

The surface soil of Buriganga accumulate various contaminants having different environmental pollutants that need urgent removal. Therefore, an attempt was taken for the evaluation of the phytoremediation potential of Indian mustard as well as rice and wheat, which are the two most important cereal crops of Bangladesh against the three important soil pollutants *viz* Pb, Cd and Cr. Moreover, Buriganga river of Dhaka city is polluted in a rapid pace and physicochemical process to remove the pollutants is not possible due to huge cost. To save the environment, the use of hyper-accumulator plants can be an effective alternative to remediate heavy metals from a polluted river, which is otherwise impossible due to lack of effort for contaminated site cleanup.

Phytoremediation is considered as a novel technology that is cheaper to clean up polluted sites in comparison to physicochemical methods (McGrath et al., 2001; Raskin et al., 1997). In this study, phytoremediation capacity of Indian mustard, rice and wheat were determined against three different heavy metals of heavily polluted Buriganga riverbank soil of Dhaka city, Bangladesh. Seeds were selected on the basis of their morphological characteristics, seed germination percentage and plant survival rate.

## MATERIALS AND METHODS

### Plant materials

Three different samples of each of Indian mustard (*Brassica juncea*), rice (*Oryza sativa*) and wheat (*Triticum aestivum*) seeds were collected from local market (Siddik bazar) of Dhaka city and the best samples were used as plant material for phytoremediation study. Selected seed samples were grown in the earthen pots in the net house of the Department of Botany of Jagannath University, Dhaka between Oct to Dec 2017. Three different samples of *Brassica juncea* seeds were labeled as B1, B2 and B3. Rice (*Oryza sativa*) and wheat (*Triticum aestivum*) seed samples were also labeled as O1, O2, O3 and T1, T2, T3 respectively (Fig 1a).

**Fig 1.**
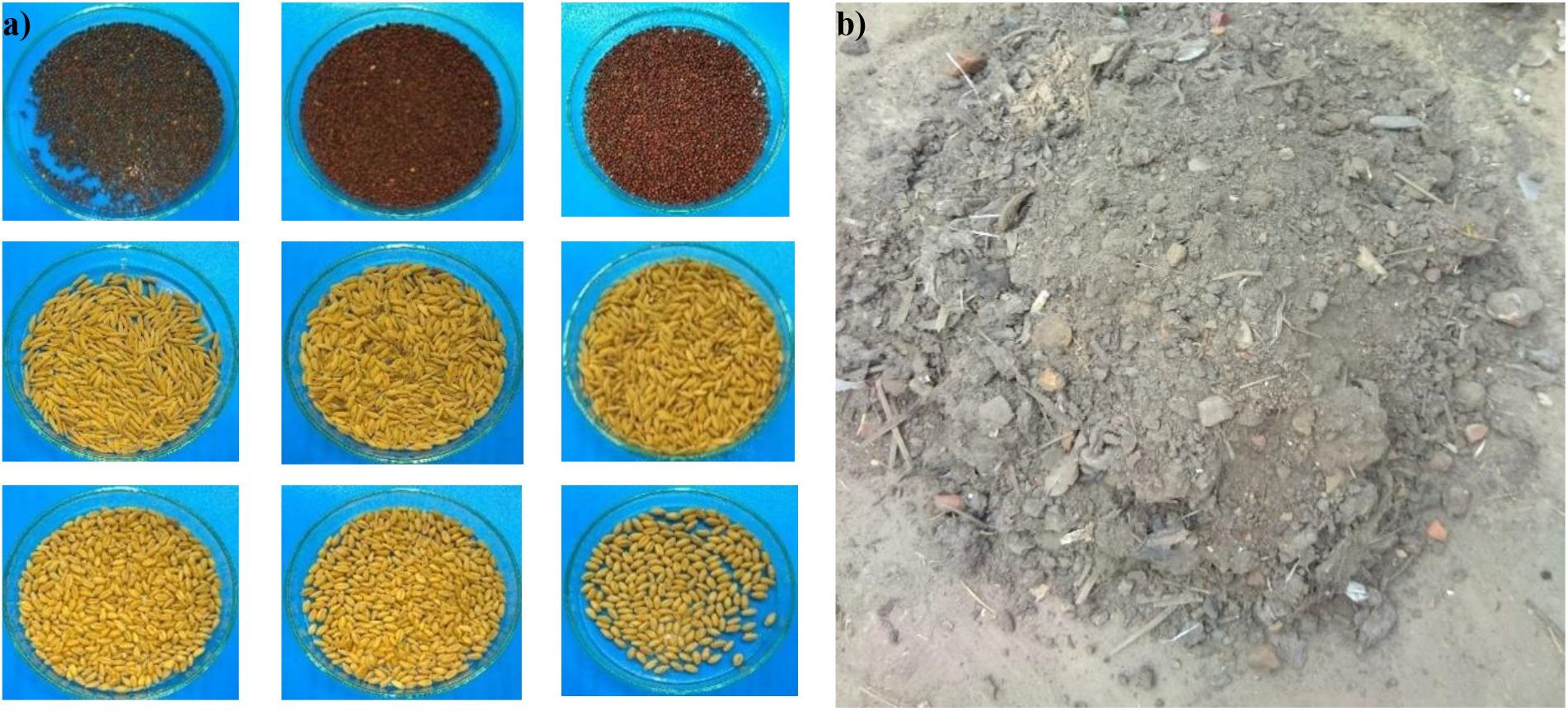
**a)** Different seed samples of *B. juncea, O. sativa* and *T. aestivum*. **b)** polluted Buriganga riverbank soil

### Source of heavy metal

Surface soil of polluted Buriganga River was used as a heavy metal source, which is the immediate uppermost layer of Buriganga riverbank containing waste materials and heavy metals (Fig 1b).

### Phytoremediation study

Heavy metals (Pb, Cr and Cd) present in the leaf and root samples of *B. juncea*, *O. sativa* and *T. aestivum* were determined at Bangladesh Council of Scientific and Industrial Research (BCSIR), Dhaka.

### Investigation of seed quality

It was very important to select the best quality of seeds to grow them under heavy metal stress condition. To do this, morphological characteristics, germination rate and plant survival rate were considered.

#### a) Morphological characteristics

Seed samples were selected for the research considering their uniform size, shape, color and disease-free appearance. Among the samples of B1, B2, B3 of *B. juncea*, the sample that showed comparatively uniform characteristics and disease-free appearance was selected for the experiment. In the same way, seeds were selected from O1, O2, O3 of *O. sativa* and T1, T2, T3 of *T. aestivum* respectively.

#### b) Seed germination percentage

To know the viability of seeds study of germination percentage is essential. Seeds with best germination percentage were selected for the experiment. Germination percentage was calculated by lab experiment. For the calculation of germination percentage seed samples were surface sterilized with 0.1% *HgCl_2_* solution for 1 minute with frequent shaking and seeds were thoroughly washed with distilled water. Samples of B1, B2, B3 of *B. juncea* were placed on the three different petri dishes containing soaked filter paper. Temperature was 20-25 °C. In the same way, samples of O1, O2, O3 of *O. sativa* and T1, T2, T3 of *T. aestivum* seeds respectively were also placed on petri dishes separately and germination percentage was observed.

Germination percentage was calculated by the following formulae-

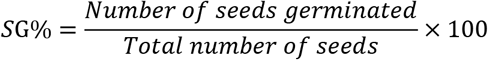

#### c) Plant survival rate

Germinating seeds of each sample were sown in different pots and calculated the plant survival rate after 15 days of sowing the seeds by the following formulae.

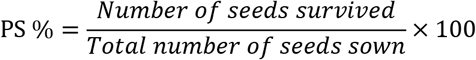

### Heavy metal treatment

#### a) Selection of seeds

The plant samples which had shown better morphological characteristics, seed germination percentage and plant survival rate were selected for heavy metal treatment. One hundred seeds of each sample were taken that was approximately 0.25g seeds of *B. juncea* and 3.00g seeds of *O. sativa* and *T. aestivum* respectively. Plant samples for phytoremediation study were collected after 30 days of seed sowing.

#### b) Soil preparation

Surface soil of polluted Buriganga riverbed was used as the source of heavy metals. Soil contains heavy metals in heterogenous condition. As a result, preparation of soil was very important for this research. Soil was mixed properly with the help of a spade. Soil containing waste material *viz*. plastic element, brickbat, small stone and some other unnecessary things were removed from the soil.

#### c) Preparation of pots

Earthen pots were used for heavy metal treatment. Three replicates were used for each plant. The pots were filled with polluted soil and each pot contained a pore at the bottom for passing extra water during and after irrigation.

#### d) Sowing of seeds

**S**eeds were grown immediately in the pots having soil with heavy metal in the net house.

### Field management

#### a) Irrigation

The plants were irrigated regularly with the help of a plastic pipe from water supply system of the net house. Usually, water was applied daily basis depending on the sun exposure.

#### b) Weed control

To avoid nutrient loss and heavy metal uptake by weeds, they were eradicated weekly. Proper care was taken during the plant growth period.

#### a) Shading

To prevent the plants from excessive sun exposure and rainwater transparent polythene bag was used to cover the top of the net house. It helped to protect the plants from being damaged by excessive heat or rainfall.

### Heavy metal tolerant plant selection

Plants of *B. juncea, O. sativa* and *T. aestivum* were selected on the basis of seed germination percentage, germination speed, vigor index, plant survival rate and hyperaccumulator group for phytoremediation study.

#### a) Seed germination rate

For the calculation of seed germination rate of all of the samples seed germination number was recorded and calculated according to the previous formulae.

#### b) Germination speed

The germination speed was calculated by the following formulae which was described by (Maguire, 1962).

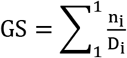

n_i_ =number of seeds germinated on i^th^ day.

D_i_=number of days from the start of the experiment.

#### c) Vigor index

Vigor index (VI) was calculated by the following formulae described by (Abdul-Baki and Anderson, 1973). Seed germination percentage (SG%) was multiplied by total seedling length (SL) and vigor index was found. Seedling length was measured on 15^th^ day of sowing the seeds.

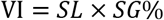

#### d) Plant survival rate

After calculating the vigor index each seedling was transferred in different pots and the plant survival rate was calculated after 30 days of seed sowing according to the previous formulae.

#### e) Plant group

To find out the hyperaccumulator plant group, plants were measured individually and according to their height they were divided into three different groups after 30 days of sowing.

### Collection of selected plant samples

Plants were collected after 30 days of seed sowing. Leaves and roots of selected plants of *B. juncea, O. sativa* and *T. aestivum* were collected and phytoremediation study was performed.

### Phytoremediation study

Pb, Cr and Cd contents in leaf and root tissues of *B. juncea*, *O. sativa* and *T. aestivum* were determined by Atomic Absorption Spectrometry (AAS).

#### a) Sample preparation

Soil was collected for the determination of metal (Pb, Cr and Cd) content. Selected plant samples of *B. juncea*, *O. sativa* and *T. aestivum* were collected from the pots of the net house and prepared for the phytoremediation study. At first, samples were washed with running tap water and then with distilled water. Next, samples were oven dried and placed on the petri dishes. Approximately 0.5g samples of *B. juncea, O. sativa* and *T. aestivum* samples were measured and placed into the beaker. 10ml conc. *HNO*_3_ (Nitric acid) was added with each sample for digestion. Samples were then placed on the hot plate. By slow boiling and evaporation process samples became concentrated. Required deionized water was added with the concentrated samples and poured into 25ml volumetric flask to prepare 25ml solution. Samples were then filtrated with Whatman no. 42 filter paper and stored in the plastic bottle for metal analysis.

#### b) Measurement of heavy metal concentration by AAS

Three concentrations of standard solution of a particular metal were selected for metal analysis. Blank solution was aspirated and adjusted to zero. A calibration curve was prepared for absorbance versus concentration of standard solution. The reading of the prepared sample solution was taken directly from the instruments. Calculation was done by the following formulae:

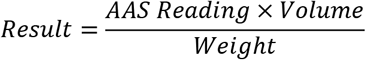

### Statistical analysis

The amount of Pb, Cd and Cr were determined using three biological replicates of each of rice, wheat and mustard samples. The mean and standard deviation were calculated.

### Experimental design

The full study was carried out by three consecutive experiments—lab experiments for observation of germination rate for seed quality selection; field experiments for selection of heavy metal tolerant plants, and again lab experiment for determination of heavy metal concentrations (Fig 2).

**Fig 2.**
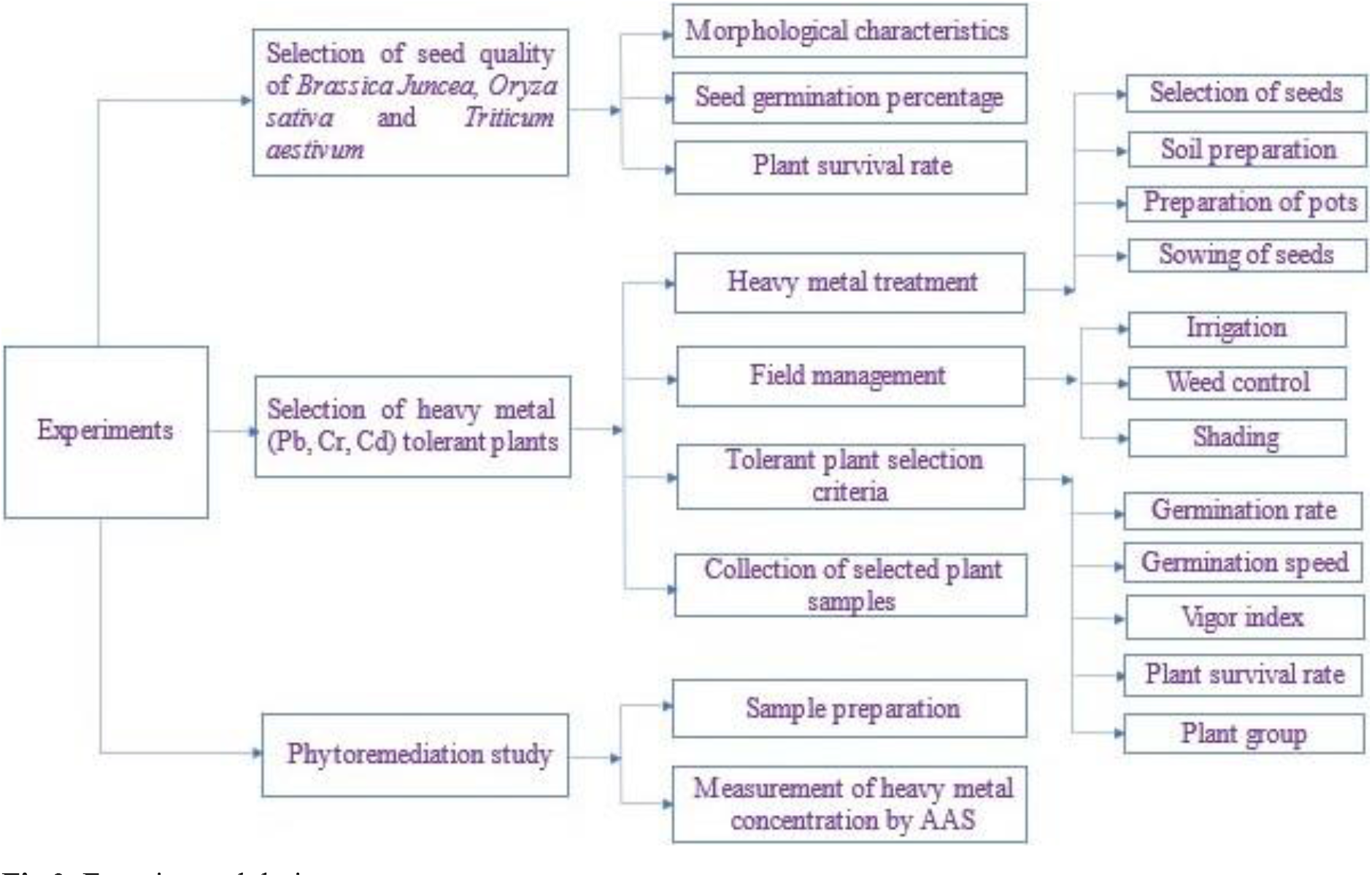
Experimental design

## RESULTS

### Morphological characteristics

Three different samples of seeds for each plant were collected from the local market to assess their quality, which is important for phytoremediation study. For this reason, best quality of seeds were selected depending on their morphological characteristics, seed germination percentage and plant survival rate (Table 1).

**Table 1.** Comparative study of different characteristics of seeds

| Sample of seeds |  | Morphological characteristics |  | Seed germination rate |  |  | Plant survival rate |  |
| --- | --- | --- | --- | --- | --- | --- | --- | --- |
|  |  | Size | Shape | Number of seeds used | Number of seeds germinated | Germination rate | Number of seeds survived | Plant survival rate |
| *Brassic juncea* | B1 | Large | Not uniform | 20 | 15 | 75% | 12 | 60% |
|  | B2 | Small | Uniform | 20 | 18 | 90% | 13 | 65% |
|  | B3 | Large | Uniform | 20 | 18 | 90% | 16 | 80% |
| *Oryza sativa* | O1 | Large | Uniform | 20 | 17 | 85% | 15 | 75% |
|  | O2 | Large | Not uniform | 20 | 16 | 80% | 12 | 60% |
|  | O3 | Small | Not uniform | 20 | 15 | 75% | 14 | 70% |
| *Triticum aestivum* | T1 | Small | Uniform | 20 | 16 | 80% | 13 | 65% |
|  | T2 | Small | Not uniform | 20 | 16 | 80% | 12 | 60% |
|  | T3 | Large | Uniform | 20 | 17 | 85% | 15 | 75% |

It was observed that size and shape are a bit different among the three species. Among the three samples (B1, B2 and B3) of *Brassica juncea* B3 showed comparatively proper size (large), shape (uniform) and colour (dark brown). Samples of *Oryza sativa* observed carefully and excellent result was found from O1 among the samples of O1, O2 and O3. O1 is comparatively large sized, uniform in shape and yellow in color. On the other hand, among the three samples of *Triticum aestivum* T3 showed the best morphological characteristics. It was comparatively large, uniform and yellow in colour (Table 1).

### Seed germination rate

Percentage of seed germination is another important parameter for seed selection. This experiment (Fig 3a) was performed in the Research Laboratory of the Department of Botany of Jagannath University, Dhaka. Percentage of seed germination was calculated after one week of seed sowing of *B. juncea*, and after two weeks of *T. aestivum* and *O. sativa* (Fig 3a) respectively.

**Fig 3.**
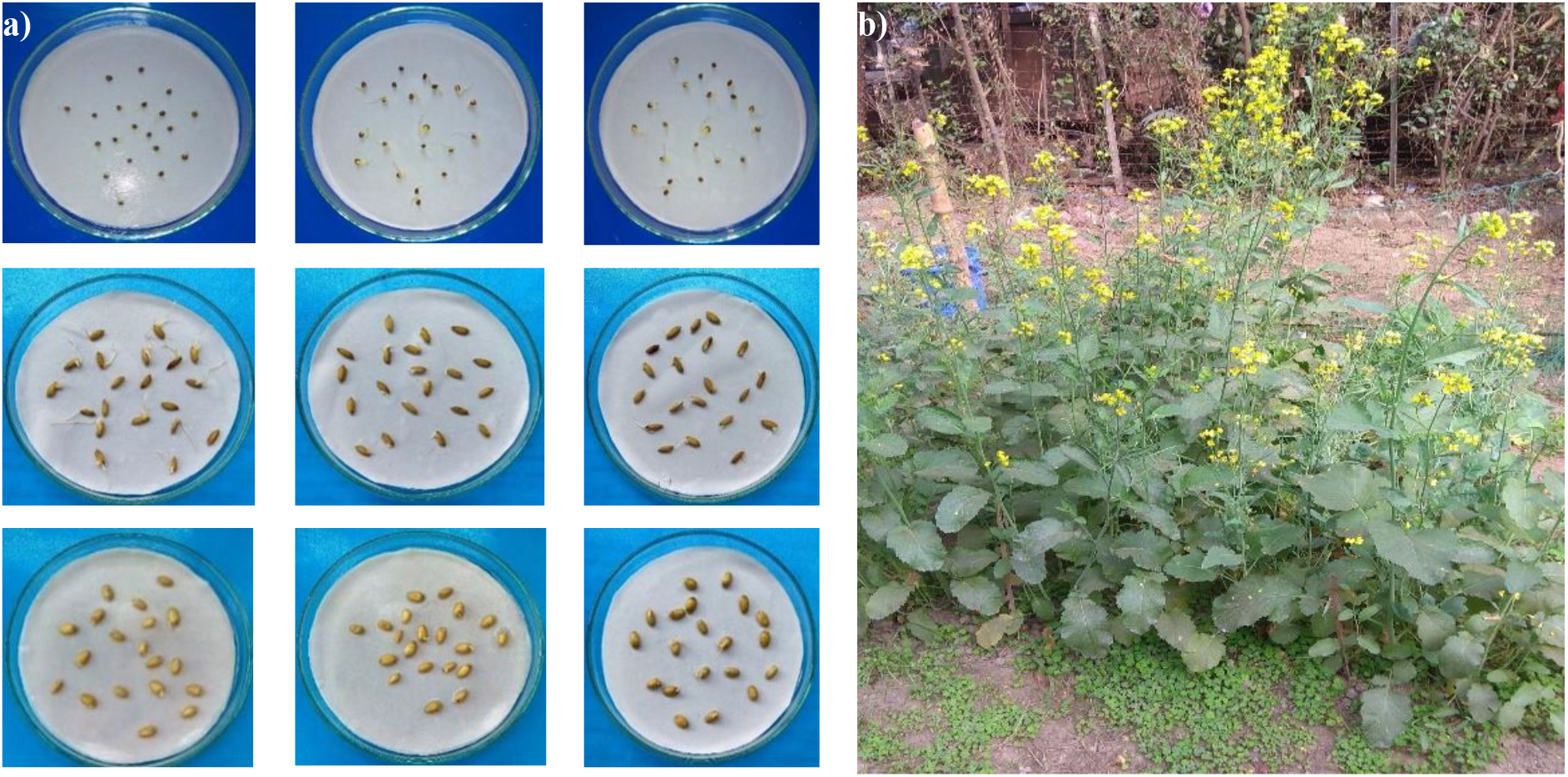
**a)** Germination of different seed samples in the lab; **b)** *B. juncea* plants in the net house

Among the three samples of *B. juncea* B2 and B3 showed equal (90%) germination percentage that was more than the sample B1 (75%). In contrast, among the three samples of *O. sativa* O1 showed the best seed germination potential. In addition, T3 sample showed the highest germination percentage among the three samples of *T. aestivum* (Table 1, Fig 3a).

### Plant survival rate

Germinated seeds were transferred in normal (uncontaminated) soil (Fig 3b) in the net house to observe their survival and plant survival data was recorded to calculate plant survival rate. Plant survival rate ranges from 60% to 80%. Among the three samples of *B. juncea* B1 showed the lowest percentage (60%) and B3 showed the highest percentage (80%). By contrast, O1 represented the highest percentage (75%) and O2 the lowest percentage (60%) of *O. sativa*. In addition, T3 gave the highest percentage (75%) and T2 the lowest (60%) of *T. aestivum*.

#### Selection of seed samples for phytoremediation study

B3 showed the best morphological character, germination percentage and plant survival rate among the three samples of *B. juncea*. As a result, B3 was selected for phytoremediation study. O1, in addition, showed the best morphological character, germination percentage and plant survival rate among the three samples of *O. sativa*. Therefore, O1 was the best sample and was selected for further analysis. T3 also showed the best morphological characteristics, seed germination percentage and plant survival rate among the three samples of *T. aestivum* and was selected for the same purpose.

### Selection of heavy metal tolerant plants

Heavy metal stress tolerant plants were selected based on five parameters—germination rate, germination speed, vigor index, plant survival rate andplant group. Selected seeds were sown in the earthen pots filled with heavy metal polluted soil in the net house for the study of these parameters.

#### Germination rate

For the calculation of seed germination rate data was recorded after one week of seed sowing of *B. juncea*, and two weeks of *T. aestivum* and *O. sativa* respectively. For the selection of hyperaccumulator genotype one hundred seeds of each type were planted out in the garden pots and 84 of *B. juncea*, 77 of *O. sativa* and 76 of *T. aestivum* were germinated. As a result, the highest germination percentage was observed in *B. juncea* (84%), followed by *O. sativa* (77%) and *T. aestivum* (76%).

#### Germination speed (GS)

After growing the seeds of *B. juncea* first seed germination was observed on the 3^rd^ day of seeds sowing and final seed germination was observed on the 5^th^ day. For the calculation of the germination speed of *O. sativa*, first seed germination was taken place on the 5^th^ day of seeds sowing and germination was completed on the 7^th^ day. Regarding *T. aestivum* first and last seed germination were observed on the 4^th^ and 6^th^ days of seed sowing (Table 2). Different seeds of *B. juncea*, *O. sativa* and *T. aestivum* were sown on the same day for observation of the germination speed. However, for the calculation of germination speed, number of observation days varied depending on the nature of germination of each sample.

**Table 2.**
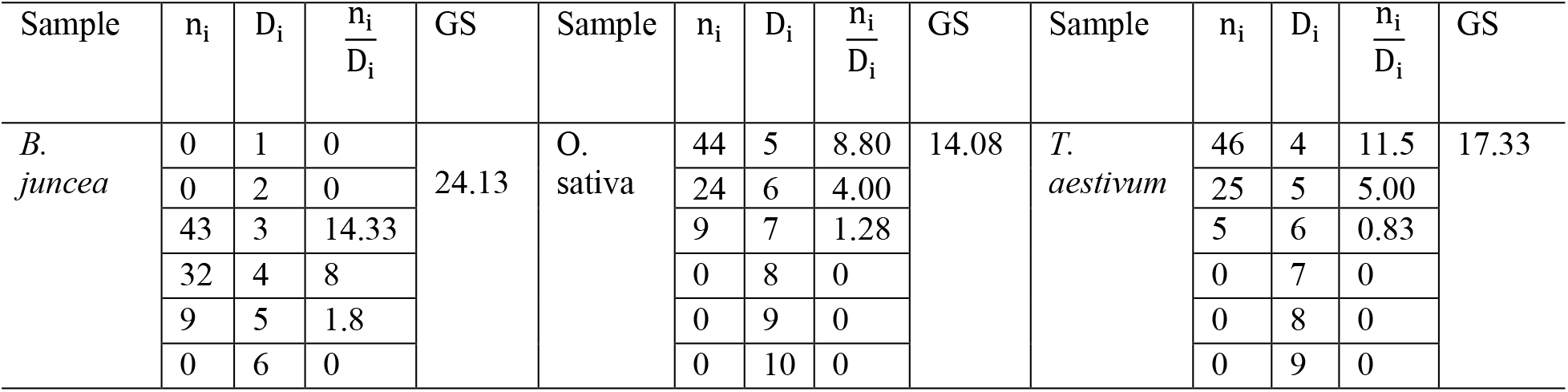
The germination speed for *Brassica juncea, Oryza sativa* and *Triticum aestivum*.

| Sample | $n_i$ | $D_i$ | $\frac{n_i}{D_i}$ | GS | Sample | $n_i$ | $D_i$ | $\frac{n_i}{D_i}$ | GS | Sample | $n_i$ | $D_i$ | $\frac{n_i}{D_i}$ | GS |
| --- | --- | --- | --- | --- | --- | --- | --- | --- | --- | --- | --- | --- | --- | --- |
| *B. juncea* | 0 | 1 | 0 | 24.13 | *O. sativa* | 44 | 5 | 8.80 | 14.08 | *T. aestivum* | 46 | 4 | 11.5 | 17.33 |
|  | 0 | 2 | 0 |  |  | 24 | 6 | 4.00 |  |  | 25 | 5 | 5.00 |  |
|  | 43 | 3 | 14.33 |  |  | 9 | 7 | 1.28 |  |  | 5 | 6 | 0.83 |  |
|  | 32 | 4 | 8 |  |  | 0 | 8 | 0 |  |  | 0 | 7 | 0 |  |
|  | 9 | 5 | 1.8 |  |  | 0 | 9 | 0 |  |  | 0 | 8 | 0 |  |
|  | 0 | 6 | 0 |  |  | 0 | 10 | 0 |  |  | 0 | 9 | 0 |  |

After calculation of germination speed, three different values were found form three different plant samples. The highest germination speed (24.13) was showed by *B. juncea* that was more than *O. sativa*. Germination speed of *T. aestivum* was higher than *O. sativa* but lower than *B. juncea*. As such, *B. juncea* is the best sample regarding the speed of germination.

#### Vigor index

The vigor index of *B. juncea* (143283.84) was higher than *O. sativa* as well as *T aestivum*. The lowest vigor index (103116.04) was recorded in *T. aestivum* (Table 3). The result of vigor index revealed that *B. juncea* is superior to *O. sativa* and *T aestivum* regarding its heavy metal tolerance.

**Table 3.** Comparative study of vigor index

| Sample | Total number of seedlings | Total length of root (cm) | Total length of Shoot (cm) | Total length of seedlings (cm) | Germination percentage (%) | Vigor index |
| --- | --- | --- | --- | --- | --- | --- |
| *B. juncea* | 84 | 439.89 | 1265.87 | 1705.76 | 84 | 143283.84 |
| *O. sativa* | 77 | 134.98 | 1589.97 | 1724.95 | 77 | 132821.15 |
| *T. aestivum* | 76 | 165.89 | 1190.90 | 1356.79 | 76 | 103116.04 |

#### Plant survival rate

Plant survival rate was calculated after 30 days of seed sowing. One hundred seeds of each species were sown and from the germinated 84, 77 and 76 plants of *B. juncea, O. sativa* and *T. aestivum*, 71, 68 and 65 plants were survived, respectively. Therefore, the highest plant survival rate for *B. juncea* is 71%, followed by *O. sativa* (68%) and *T. aestivum* (65%). This result indicates that *B. juncea*, *O. sativa* and *T. aestivum* have the ability to cope with heavy metal stress environment.

#### Plant group

Plant group with maximum survived plants according to their height was considered as one of the criteria to be considered as hyperaccumulator plant. The reason behind this is that heavy metals have toxic effect by reducing the root length as well as shoot height (Mahmood et al., 2007) and hence growth and survival of plants. The group which contained more plants is the indication that the plant group is capable of coping with heavy metal stress condition. As such, plants were divided into three groups according to their height. The survived 71 plants of *B. juncea* was divided into short (15-25 cm), medium (26-35 cm) and long (36-45 cm) plant groups and the highest number (41) of plants were survived in the medium plant group followed by long (19) and short (11) plant groups. In the same way, the survived 68 plants of *O. sativa* were divided into short (25-35 cm), medium (36-45 cm) and long (46-55 cm) plant groups and long plant group has the highest number (42) of survived plants followed by medium (15) and short (11) groups. There were 65 survived plants of *T. aestivum*, which were also subdivided into short (15-25 cm), medium (26-35 cm) and long (36-45 cm) plant groups, of which medium plant group has the highest number (30) of survived plants followed by short (20) and long (15) groups. Heavy metal tolerant three plants were selected randomly from the highest plant surviving groups of each plant for the assessment of phytoremediation.

### Assessment of phytoremediation potential

Pb, Cr and Cd contents in the root and leaf samples of *B. juncea, O. sativa* and *T. aestivum* were determined by Atomic Absorption Spectrometry (AAS). The selected plants were planted in the earthen pot filled with polluted soil with heavy metal (Fig 4).

**Fig 4.**
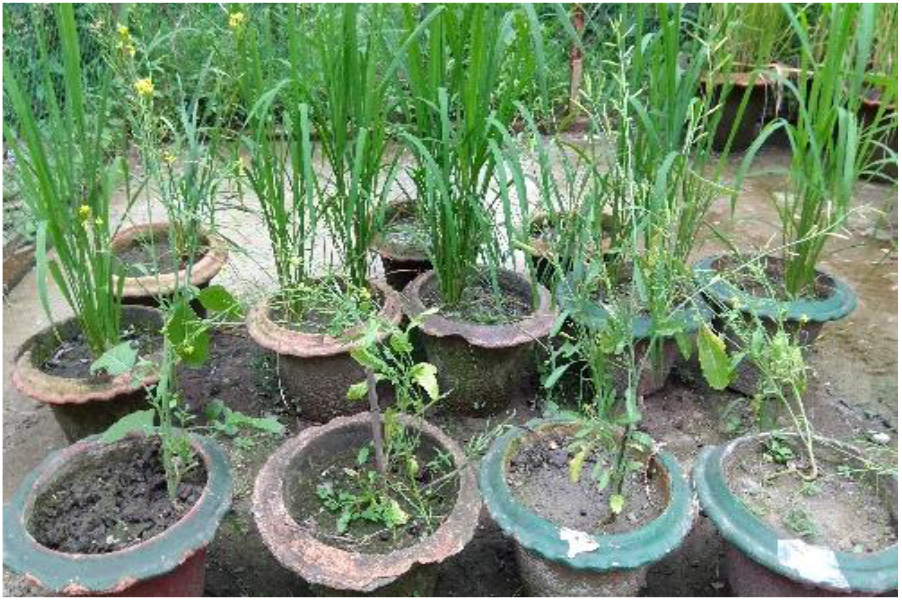
*B. juncea*, *O. sativa* and *T. aestivum* plants growing in the garden pot under heavy metal stress condition.

#### Determination of Pb content

Phytoremediation study was conducted using heavy metal polluted surface soil of Buriganga river and heavy metal accumulation/removal capacity of leaf and root samples of *B. juncea, O. sativa* and *T. aestivum*. Amount of Pb, Cr and Cd present in different samples were determined and results were recorded (Table 4). It was observed that the highest amount (11.6755±1.9860 mg/kg dry weight) of Pb was found in the leaf sample of *B. juncea* that was approximately three times greater than *T. aestivum* and more than the value of *O. sativa*. On the other hand, the highest amount (51.4251±5.0320 mg/kg dry weight) of Pb was found in the root sample of *B. juncea* that was greater than *O. sativa* and *T. aestivum* respectively.

**Table 4.**
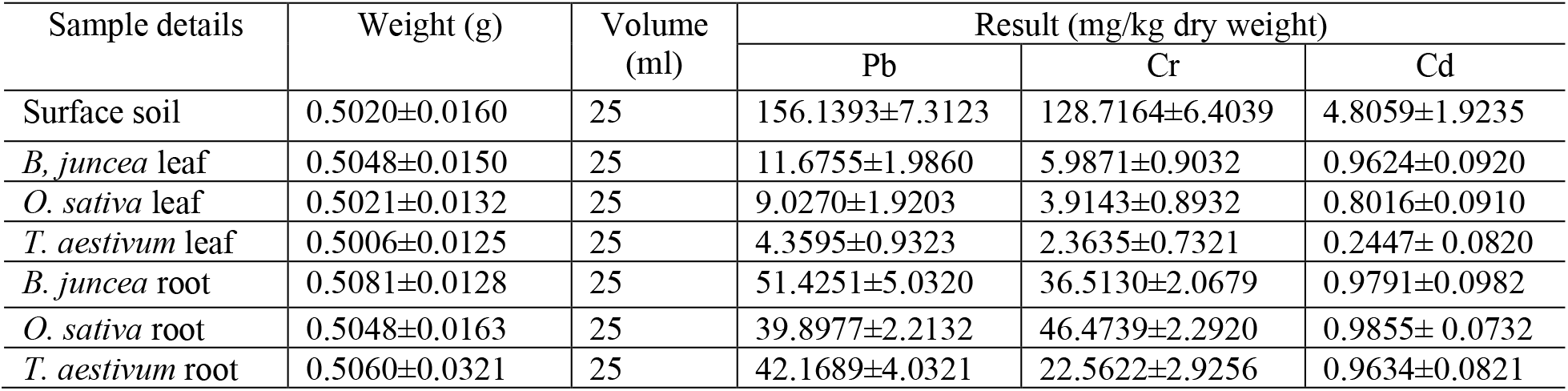
Comparative amount of Pb, Cr, Cd in the leaf and root of *Brassica juncea, Oryza sativa* and *Triticum aestivum* (n=3, ±= standard deviation).

The result of accumulation of Pb in the leaf and root of *B. juncea* found by (Choudhury et al., 2016) is 5.6 and 5.85 mg/kg dry weight respectively and is two- and ten-times less than the concentration of Pb in the leaf and root of current study. Concentration of Pb in the leaf and root of paddy rice of earlier research (Payus and Talip, 2014) was 0.26 ± 0.14 and 7.70 ± 1.27 mg/kg respectively, which are 35- and 5-fold less than the current research. Pb concentration in the root of wheat (18.49–30.83 mg/kg) found by (Liu et al., 2009) was less than the result found in this research.

#### Determination of Cr content

The highest amount (5.9871±0.9032 mg/kg dry weight) of Cr was found in the leaf samples of *B. juncea* that was approximately two-fold higher than *T. aestivum* and more than *O. sativa*. In contrast, the highest amount (46.4739± 2.2920 mg/kg dry weight) of Cr was found in the root samples of *O. sativa* that was approximately 2 times greater than *T. aestivum* and more than *B. juncea* (Table 4).

Cr concentration in the leaf of *B. juncea* of arlier research (Choudhury et al., 2016) of is 8.4 mg/kg, which is more than the current research; and in the root it is 20.1 mg/kg, which is less than the present study. On the other hand, previous research (Payus and Talip, 2014) found higher Cr concentration of rice plant in the leaf (4.34 ± 2.01 mg/kg) and root (5.46 ± 2.26 mg/kg) samples respectively, than the result obtained by present study. From the research of (Liu et al., 2009), Cr concentration found in the root (2.55–8.07 mg/kg) of wheat was less than the result of current research.

#### Determination of Cd content

Cd accumulation in the leaf samples of *B. juncea* was 0.9624±0.0920 mg/kgdry weight) followed by *O. sativa*. However, the lowest amount (0.2447± 0.0820 mg/kg dry weight) was found in the leaf of *T. aestivum*. Which is more than three times lower than *B. juncea* and *O. sativa* leaves. In the root samples, the amount of Cd was approximately same in all samples (Table 4).

Cd concentration by earlier research (John et al., 2009) in the root of *B. juncea* is 0.32±0.02mg/kgat 25 days, which is less than the result obtained by present study. Another research (Payus and Talip, 2014) found Cd concentration in the leaf and root of rice as 0.11 ± 0.05 and 0.38 ± 0.09 mg/kg respectively, which are seven and three times less than the present study. From the research of (Liu et al., 2009) Cd concentration of root (2.63–4.83mg/kg) of *T. aestivum* was more than the result of this research.

## DISCUSSION

Environment is polluted by different waste materials and are the most important fountain of heavy metals which have harmful effect on both water and soil. In addition, heavy metals have various inhibitory effect on the growth and metabolism of plants (Boussama et al., 1999; John et al., 2009). The removal of heavy metals by hyperaccumulator plants is 100 to 1,000-fold more compared to the non-accumulator species (Tangahu et al., 2011). Different methods of phytoremediation are applied for wastewater treatment and removal of soil pollutants (Dordio and Carvalho, 2013). Phytoremediation method is one of the most ecofriendly method than physico-chemical techniques (Salido et al., 2003).

In the present study, *B. juncea*, *O. sativa* and *T. aestivum* were used to assess their phytoremediation potential against three most important environmental pollutants (Pb, Cd and Cr) of polluted Buriganga riverbank soil of Dhaka city, Bangladesh. Leaf and root of each plant were used to know their capacity to remove the contaminants as hyperaccumulators.

During the selection of seed samples, it was also observed that the highest germination percentage (90%) and plant survival rate (80%) was found in the B3 sample of *B. juncea*. The equal percentage of seed germination (85%) and plant survival rate (75%) was recorded in the sample of O1 of *O. sativa* and T3 of *T. aestivum*, respectively. Based on the morphological characteristics, percentage of seed germination and plant survival rate B3, O1 and T3 samples were selected from *B. juncea*, *O. sativa* and *T. aestivum* respectively for phytoremediation study. The result also revealed that regarding the aforementioned criteria *B. juncea* is one of the best experimental materials for the study of phytoremediation.

To select the best genotypes for heavy metal phytoremediation, seed germination rate, germination speed, vigor index, plant survival rate and plant group were considered. From one hundred seeds of each plant sown in the earthen pots with heavy metal polluted soils, germination rate of *B. juncea, O. sativa* and *T. aestivum* was 84%, 77% and 76% respectively. In addition, seed germination speed of *B. juncea, O. sativa* and *T. aestivum* was found on the 3^rd^, 5^th^ and 4^th^ day of seed sowing completed on 5^th^, 7^th^ and 6^th^ day respectively. On the other hand, the highest vigor index was found in *B. juncea* followed by *O. sativa* and *T. aestivum*. Same trend was observed regarding the plant survival rate. From these results it can be concluded that *B. juncea*, *O. sativa* and *T. aestivum* are capable of coping with heavy metal stress condition; however, *B. juncea* is the best heavy metal tolerant plant regarding these parameters. Finally, three plants were randomly selected from the highest plant surviving groups of *B. juncea, O. sativa* and *T. aestivum*, respectively for assessment of phytoremediation.

It was found that root samples contained the highest amount of three types of heavy metals. The highest amount (51.4251± 5.0320 mg/kg dry weight) of Pb that was found in the root samples of *B. juncea* is approximately 5 times greater than leaf samples (11.6755± 1.9860 mg/kgdry weight) of *B. juncea*. By contrast, the highest amount (42.1689± 4.0321mg/kg dry weight) of Pb found in root samples of *T. aestivum* is approximately 10 times greater than leaf samples (11.6755± 1.9860 mg/kg dry weight) of *T. aestivum*. Similar trend was obtained regarding the leaf and root samples of *O. sativa*.

In summary, it is revealed that in comparison to *O. sativa* and *T. aestivum, B. juncea* is the highest hyperaccumulator species regarding Pb, Cr and Cd accumulation in the leaf and root tissues (Table 4). Moreover, from these experiments it is revealed that *B. juncea*, *O. sativa* and *T. aestivum* have the ability to produce hyperaccumulator genotypes. It is also alarming that not only the root uptakes heavy metal but also the leaf accumulates heavy metal during the heavy metal stress condition. As a result, it may be concluded that if the plants are used in phytoremediation study, then it must be assured that the edible plant parts must not be used as food or fodder.

## Acknowledgement

The authors are grateful to Mohammad Moniruzzaman, Senior Scientific Officer, Bangladesh Council of Scientific and Industrial Research (BCSIR), Dhaka, Bangladesh for his help in conducting phytoremediation study. The authors also show their gratitude to Jagannath University for project assistance through the University Grant Commission of Bangladesh.

## References

Abdul-Baki, A. A., & Anderson, J. D. (1973). Vigor Determination in Soybean Seed by Multiple Criteria 1. Crop Science, 13(6), 630–633. doi:10.2135/cropsci1973.0011183X001300060013x

Ahmad, P., Nabi, G., & Ashraf, M. (2011). Cadmium-induced oxidative damage in mustard [ Brassica juncea (L.) Czern. & Coss.] plants can be alleviated by salicylic acid. South African journal of botany, 77(1), 36–44. doi:10.1016/j.sajb.2010.05.003

Ampiah-Bonney, R. J., Tyson, J. F., & Lanza, G. R. (2007). Phytoextraction of Arsenic from Soil by Leersia Oryzoides. Int J Phytoremediation, 9(1), 31–40. doi:10.1080/15226510601139383

Boussama, N., Ouariti, O., Suzuki, A., & Ghorbal, M. H. (1999). Cd-Stress on Nitrogen Assimilation. Journal of plant physiology, 155(3), 310–317. doi:10.1016/S0176-1617(99)80110-2

Chandra, R., Bharagava, R. N., Yadav, S., & Mohan, D. (2009). Accumulation and distribution of toxic metals in wheat (Triticum aestivum L.) and Indian mustard (Brassica campestris L.) irrigated with distillery and tannery effluents. J Hazard Mater, 162(2), 1514–1521. doi:10.1016/j.jhazmat.2008.06.040

Choudhury, M. R., Islam, M. S., Ahmed, Z. U., & Nayar, F. (2016). Phytoremediation of heavy metal contaminated buriganga riverbed sediment by Indian mustard and marigold plants. Environ. Prog. Sustainable Energy, 35(1), 117–124. doi:10.1002/ep.12213

Dordio, A. V., & Carvalho, A. J. P. (2013). Organic xenobiotics removal in constructed wetlands, with emphasis on the importance of the support matrix. J Hazard Mater, 252-253, 272–292. doi:10.1016/j.jhazmat.2013.03.008

Garbisu, C., & Alkorta, I. (2001). Phytoextraction: a cost-effective plant-based technology for the removal of metals from the environment. Bioresour Technol, 77(3), 229–236. doi:10.1016/S0960-8524(00)00108-5

Gaur, A. a. A., A. (2004). Prospects of arbuscular mycorrhizal fungi in phytoremediation of heavy metal contaminated soils. Current Science, 86(4).

Hussain, B., Ashraf, M. N., Shafeeq ur, R., Abbas, A., Li, J., & Farooq, M. (2021). Cadmium stress in paddy fields: Effects of soil conditions and remediation strategies. Sci Total Environ, 754, 142188. doi:10.1016/j.scitotenv.2020.142188

Ishikawa, S., Ae, N., Murakami, M., & Wagatsuma, T. (2006). Is Brassica juncea a suitable plant for phytoremediation of cadmium in soils with moderately low cadmium contamination?: Possibility of using other plant species for Cd-phytoextraction. Soil science and plant nutrition (Tokyo), 52(1), 32–42. doi:10.1111/j.1747-0765.2006.00008.x

John, R., Ahmad, P., Gadgil, K., & Sharma, S. (2009). Heavy metal toxicity: Effect on plant growth, biochemical parameters and metal accumulation by Brassica juncea L. International journal of plant production, 3(3), 65–75.

Liu, W. X., Liu, J. W., Wu, M. Z., Li, Y., Zhao, Y., & Li, S. R. (2009). Accumulation and Translocation of Toxic Heavy Metals in Winter Wheat (Triticum aestivum L.) Growing in Agricultural Soil of Zhengzhou, China. Bull Environ Contam Toxicol, 82(3), 343–347. doi:10.1007/s00128-008-9575-6

Maguire, J. D. (1962). Speed of Germination—Aid In Selection And Evaluation for Seedling Emergence And Vigor 1. Crop Science, 2(2), 176–177. doi:10.2135/cropsci1962.0011183X000200020033x

Mahmood, T., Islam, K. R., & Muhammad, S. (2007). Toxic effects of heavy metals on early growth and tolerance of cereal crops. Pakistan journal of botany, 39(2), 451–462.

Maine, M. A., Hadad, H. R., Camano Silvestrini, N. E., Nocetti, E., Sanchez, G. C., & Campagnoli, M. A. (2021). Cr, Ni, and Zn removal from landfill leachate using vertical flow wetlands planted with Typha domingensis and Canna indica. Int J Phytoremediation, 1–10. doi:10.1080/15226514.2021.1926909

McGrath, S. P., Zhao, F. J., & Lombi, E. (2001). Plant and rhizosphere processes involved in phytoremediation of metal-contaminated soils. Plant and soil, 232(1/2), 207–214. doi:10.1023/A:1010358708525

McIntyre, T. (2003). Phytoremediation. In (pp. 97–123). Berlin, Heidelberg: Springer Berlin Heidelberg.

McIntyre, T., & Lewis, G. M. (1997). The advancement of phytoremediation as an innovative environmental technology for stabilization, remediation, or restoration of contaminated sites in Canada: A discussion paper. Journal of soil contamination, 6(3), 227–241.

Melo, G. W., Furini, G., Brunetto, G., Comin, J. J., Simão, D. G., Marques, A. C. R., … Zalamena, J. (2021). Identification and phytoremediation potential of spontaneous species in vineyard soils contaminated with copper. International Journal of Phytoremediation, 1–8. doi:10.1080/15226514.2021.1940835

Murakami, M., Nakagawa, F., Ae, N., Ito, M., & Arao, T. (2009). Phytoextraction by Rice Capable of Accumulating Cd at High Levels: Reduction of Cd Content of Rice Grain. Environ. Sci. Technol, 43(15), 5878–5883. doi: 10.1021/es8036687

Mwegoha. (2008). The use of phytoremediation.

Payus, C., & Talip, A. F. A. (2014). Assessment of heavy metals accumulation in paddy rice (Oryza sativa). African journal of agricultural research, 9(41), 3082–3090. doi:10.5897/AJAR2014.8760

Prasad, M. N. V., & Freitas, H. M. D. (2003). Metal hyperaccumulation in plants - Biodiversity prospecting for phytoremediation technology. Electron. J. Biotechnol, 6(3), 285–321. doi:10.4067/S0717-34582003000300012

Pratas, J., Favas, P. J. C., Paulo, C., Rodrigues, N., & Prasad, M. N. V. (2012). URANIUM ACCUMULATION BY AQUATIC PLANTS FROM URANIUM-CONTAMINATED WATER IN CENTRAL PORTUGAL. Int J Phytoremediation, 14(3), 221–234. doi:10.1080/15226514.2011.587849

Raskin, I., Smith, R. D., & Salt, D. E. (1997). Phytoremediation of metals: using plants to remove pollutants from the environment. Curr Opin Biotechnol, 8(2), 221–226. doi:10.1016/S0958-1669(97)80106-1

Rodriguez, L., Lopez-Bellido, F. J., Carnicer, A., Recreo, F., Tallos, A., & Monteagudo, J. M. (2005). Environmental Chemistry. In (pp. 197–204). Berlin, Heidelberg: Springer Berlin Heidelberg.

Salido, A. L., Hasty, K. L., Lim, J.-M., & Butcher, D. J. (2003). Phytoremediation of Arsenic and Lead in Contaminated Soil Using Chinese Brake Ferns (Pteris vittata) and Indian Mustard (Brassica juncea). Int J Phytoremediation, 5(2), 89–103. doi:10.1080/713610173

Satpathy, D., Reddy, M. V., & Dhal, S. P. (2014). Risk Assessment of Heavy Metals Contamination in Paddy Soil, Plants, and Grains (Oryza sativa L.) at the East Coast of India. Biomed Res Int, 2014, 545473–545411. doi:10.1155/2014/545473

Shukla, G. S., & Singhal, R. L. (1984). The present status of biological effects of toxic metals in the environment: lead, cadmium, and manganese. Revue canadienne de physiologie et pharmacologie, 62(8), 1015–1031. doi:10.1139/y84-171

Tangahu, B. V., Siti Rozaimah Sheikh, A., Hassan, B., Mushrifah, I., Nurina, A., & Muhammad, M. (2011). A Review on Heavy Metals (As, Pb, and Hg) Uptake by Plants through Phytoremediation. International Journal of Chemical Engineering, 2011. doi:10.1155/2011/939161

Wan, X., Lei, M., & Chen, T. (2016). Cost–benefit calculation of phytoremediation technology for heavy-metal-contaminated soil. Sci Total Environ, 563-564, 796–802. doi:10.1016/j.scitotenv.2015.12.080

Wang, Q.-R., Liu, X.-M., Cui, Y.-S., Dong, Y.-T., & Christie, P. (2002). RESPONSES OF LEGUME AND NON-LEGUME CROP SPECIES TO HEAVY METALS IN SOILS WITH MULTIPLE METAL CONTAMINATION. J Environ Sci Health A Tox Hazard Subst Environ Eng, 37(4), 611–621. doi:10.1081/ESE-120003241

Zakaria, Z., Zulkafflee, N. S., Mohd Redzuan, N. A., Selamat, J., Ismail, M. R., Praveena, S. M., … Abdull Razis, A. F. (2021). Understanding potential heavy metal contamination, absorption, translocation and accumulation in rice and human health risks. Plants (Basel), 10(6), 1070. doi:10.3390/plants10061070

